# Rapid Genotype Refinement for Whole-Genome Sequencing Data using Multi-Variate Normal Distributions

**DOI:** 10.1101/031484

**Authors:** Rudy Arthur, Jared O’Connell, Ole Schulz-Trieglaff, Anthony J. Cox

## Abstract

Whole-genome low-coverage sequencing has been combined with linkage-disequilibrium (LD) based genotype refinement to accurately and cost-effectively infer genotypes in large cohorts of individuals. Most genotype refinement methods are based on hidden Markov models, which are accurate but computationally expensive. We introduce an algorithm that models LD using a simple multivariate Gaussian distribution. The key feature of our algorithm is its speed, it is hundreds of times faster than other methods on the same data set and its scaling behaviour is linear in the number of samples. We demonstrate the performance of the method on both low-coverage and high-coverage samples.

**Availability**: The source code is available at https://github.com/sequencing/marvin

## 1 INTRODUCTION

The 1000 Genomes Project (1000GP) has pioneered the approach of combining low-coverage whole-genome sequencing (LCWGS) with LD-based genotype refinement to successfully build large panels of accurately genotyped individuals (The 1000 Genomes Project Consortium, 2010, 2012, 2015). This has provided a cost-effective alternative to sequencing many individuals at high-coverage. However, genotype refinement has a large computational burden. For example, Delaneau *et al*. (2014) quote around 32 compute years to perform haplotype estimation on 1 092 LCWGS individuals using the 1000GP haplotype estimation pipeline. Given increasing sample sizes, decreasing sequencing costs and the typically super-linear scaling of refinement algorithms, we are fast approaching a point where computation will account for a substantial proportion of the cost of such analyses.

Low-coverage genotyping typically proceeds by calculating genotype likelihoods (GLs) at a fixed set of variants (SNPs and small indels) from read alignments, the variant list being created at an earlier variant discovery step. These GLs reflect the likelihood of the read data conditional on each of the three possible genotypes (assuming a bi-allelic site). These uncertain GLs are then refined into genotypes by exploiting linkage disequilibrium (LD), the correlation between physically close variants across individuals. This final step is often referred to as *genotype refinement* and involves one (or more) phasing and imputation algorithms. The most accurate phasing and imputation techniques typically employ hidden Markov models (HMMs) which are computationally demanding, examples include Beagle (Browning and Browning, 2007), Thunder (Li *et al*., 2011) and SHAPEIT (Delaneau *et al*., 2012, 2013). The final genotypes of 1000GP were created using a combination of SHAPEIT and Beagle; starting haplotypes where generated with the faster Beagle method and then were further refined using the slower, and more accurate, SHAPEIT (Delaneau *et al*., 2014).

A closely related problem is the imputation of variants into study samples assayed on DNA microarrays from reference panels of sequenced individuals (Marchini *et al*., 2007). Several very fast methods have recently emerged for this scenario (Howie *et al*., 2012; Durbin, 2014; Fuchsberger *et al*., 2015). These rely on the availability of phased haplotypes for both study and reference data and it is not clear such algorithms will generalise to the LCWGS use case.

An alternative to HMM-based imputation is simply to predict genotypes as linear combinations of other genotypes at physically close flanking markers, modelling the correlation between variants as a multivariate normal (MVN) distribution. This idea was first introduced by Wen and Stephens (2010), where it was used in the more traditional setting of imputing genotypes into DNA microarray samples from a reference panel. Menelaou and Marchini (2013) introduced a related approach, MVNcall, that performs imputation on LCWGS data for which the individual has also been assayed on a DNA microarray, exploiting the “backbone” of confident microarray genotypes to improve genotypes at non-microarray sites.

We introduce a new technique based on MVN representations of linkage disequilibrium that extends these ideas to the LCWGS-only imputation scenario. The method exploits various efficient linear algebra operations, making it hundreds of times faster than the fastest HMM method. This speed comes with a decrease in accuracy compared to HMMs, but is still substantially more accurate than genotype calls made using no LD information.

In the methods section we outline the model and its implementation. In our results section we contrast the speed and accuracy of our technique with Beagle on 2 535 samples from 1000GP Phase 3 (LCWGS) and 3 781 samples taken from the UK10K project (UK10K Consortium *et al*., 2015; Huang *et al*., 2015). Finally, we demonstrate the applicability of LD-based genotype refinement in the high-coverage WGS setting, something that has not been investigated to date.

The method is implemented in a software package called MarViN (MultiVariate Normal imputation) and is freely available under the GPLv3 license.

## 2 MATERIALS AND METHODS

We assume that *N* diploid individuals have been sequenced and used to detect *M* bi-allelic polymorphisms. We record the number of copies of the non-reference (alternate) allele in a matrix

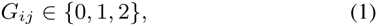
where the indexes *i* and *j* label polymorphic sites and individuals respectively. We assume that we have been given genotype likelihoods (GLs)

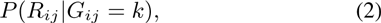
where *k* ∈ {0, 1, 2} and *R_ij_* denotes the reads aligning to site *i* in individual *j*.

### 2.1 Single-site model

We now describe a simple Expectation-Maximization (EM) algorithm that we use to initialise our model. We apply Bayes’ theorem to obtain posterior probabilities of genotypes:

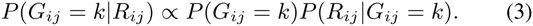
where *P*(*G_ij_ = k*) is the prior probability of seeing genotype *k* and is initialised as 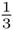. Dos can be calculated from

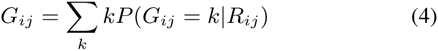

This constitutes the E-step of our routine. The M-step involves re-estimating our prior, *P*(*G_ij_* = *k*). First, we estimate site allele frequencies as

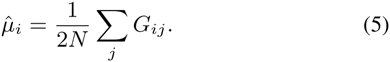

Assuming Hardy-Weinberg equilibrium, our updated prior is then

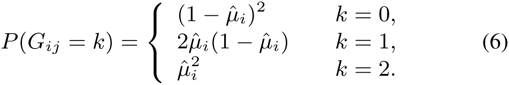

The E-step and M-step are iterated and generally converge rapidly.

### 2.2 Multi-site model

This EM algorithm gives an estimate of *G_ij_* that takes into account the population allele frequency at site *i* but ignores any correlation with flanking sites (*i.e*. linkage disequilibrium). We now describe how to improve the estimate of *G* using LD. A simple way to encode LD is with the *M × M* covariance matrix Σ, where

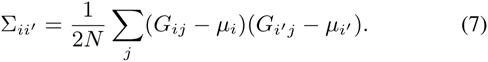

Following Wen and Stephens (2010), we make the assumption that the probability density for the vector of dosages *g*^(^*^j^*^)^ for individual *j*, the *j*^th^ column of the genotype matrix 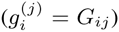, is multivariate normal (MVN):

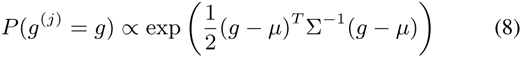

We can then ask “what is the distribution for the dosage at site *i* of individual *j* conditional on the dosages at all other sites?” For the MVN, a closed form expression for this conditional probability exists:

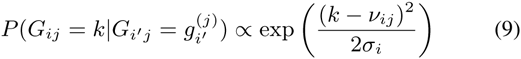
where 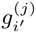 refers to all genotypes excluding site *i* and

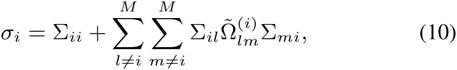

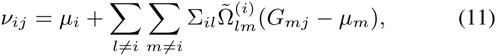
and the matrix 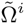 is the inverse of the matrix formed by deleting the *i*^th^ row and column from Σ.

In words, what we are doing is using the genotype matrix to estimate allele frequencies and linkage disequilibrium. Fixing these, we re-estimate the genotype matrix using the MVN assumption. The approach is similar to our single site EM algorithm, but with the simple population frequency prior in equation 6 replaced with the more sophisticated population LD prior in equation 9.

Examining the terms closely, we see that *σ_i_* is independent of the individual, as is the quantity

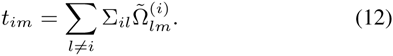

Thus we need only calculate it once. Rewriting equation 11, we see that updating the mean of individual *j* is achieved by evaluating

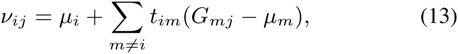
at a cost of one dot product per site per individual.

### 2.3 Algorithm description

We initialise *G* using the single-site model described in Section 2.1.

We then repeat the following steps for a default of five iterations:

1. Calculate *μ* and Σ from *G*.
2. Calculate *t_im_* and *σ_i_* for all sites *i* and *m*.
3. For all sites *i* and individuals *j*, calculate *v_ij_*.
4. Update *P*(*G_ij_* = *k|R_ij_*) using equations 3 and 9.
5. Recalculate *G*.

We take the final estimate of *G* as our imputed genotypes. We could iterate steps 2 and 3, reusing the covariance matrix obtained at the beginning of the iteration but we found this to be unhelpful in practice.

### 2.4 Calculating Ω

Computationally, Step 1 is dominated by the calculation of Σ, which takes *O*(*NM*^2^) operations. Step 2 requires a matrix vector product for every individual and so is also *O*(*NM*^2^). However, a straight-forward implementation of Step 3 would be *O*(*M*^4^), since a matrix must be inverted at each site at a cost of *O*(*M*^3^) per inversion.

To see how Step 3 can be sped up, consider the case where we want to update the marker 1 while fixing the M − 1 markers to the right. We write the covariance matrix in the following form:

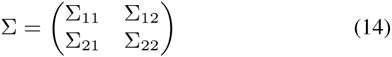
where Σ_11_ is 1 × 1 and Σ_22_ is (*M* − 1) × (*M* − 1).

To calculate *σ*_1_, we require

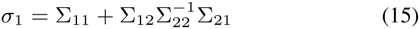
(compare equation 10). The big overhead here is calculating 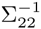. We define

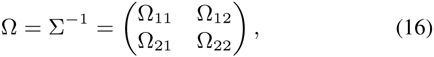
where the blocks are sized to match the corresponding submatrices of Σ. By making an LDU decomposition, we can show that

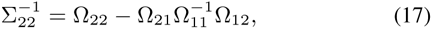
which is known as the Schur complement of Ω_11_ in Ω. This gives us 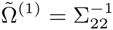 which we can use in equation 12.

Consider the variant at site *i*. The matrix we need to invert in order to evaluate the conditional expectation is the inverse of a submatrix of Σ formed by deleting the *i*^th^ row and column of Σ. Swapping rows *i* and 1 and columns *i* and 1 of Σ puts the matrix we need the inverse of in the position of Σ_22_ in equation 14. A row and column can be swapped by pre and post multiplying with a permutation matrix *P*.

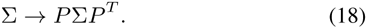

Because permutation matrices are orthogonal we have that

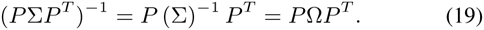

The required inverse for variant i can be obtained by applying equation 17 again on the permuted matrix. In practice we just swap rows and columns of the matrix the usual way, which is equivalent to the multiplication. This trades *M* matrix inverses for a single matrix inverse plus *M* matrix operations of complexity *O*(*M*^2^) each (matrix-vector products), giving an *O*(*M*^3^) overall cost.

### 2.5 Using a reference panel

If we have a small number of individuals to impute and a reference panel formed from a large number of individuals with hard genotypes assigned, we can impute individuals using the panel by following the procedure below:

1. Calculate allele frequencies *μ* and the covariance matrix Σ from the panel.
2. Use the panel allele frequencies to obtain an initial estimate of G from the GLs.
3. Calculate *t_im_* and *σ_i_* for all sites *i*.
4. For each individual with genotype *g*^(^*^j^*^)^ to be imputed, the following steps are performed *K* times:

1. For all sites *i*, calculate *v_ij_*
2. Update 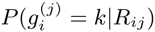 using Bayes’ theorem, equations 3 and 9.
3. Recalculate *g*^(^*^j^*^)^.

Calculating *t_im_* is *O*(*M*^3^) and Σ is *O*(*NM*^2^), both of which must be done once per panel. To impute each new individual then requires performing *O*(*M*^2^) operations for each of *K* iterations, where *K* will be around 5 in practice.

### 2.6 Regularizing the covariance matrix

To guard against degeneracy due to perfect correlation and force the variance to be non-zero, we performed Tikhonov regularization on the covariance matrix, *i.e*. applied the transformation

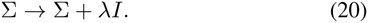

We found 0.06 to be an effective value for the regularization parameter λ in all of our tests. Alternative regularization methods (such as adding a matrix proportional to the diagonal of the covariance matrix, as done in the Levenberg-Marquardt algorithm) were evaluated but were not found to confer a significant improvement.

After Wen and Stephens (2010), we also modify the mean as follows:

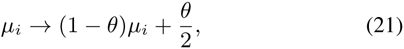
where

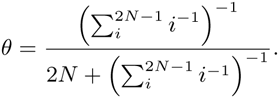

This correction is relevant in the case of small cohorts where the empirical mean may be a bad estimate of the true mean, the specific form above is derived in Wen and Stephens (2010) using the model of Li and Stephens (2003). In our case, with cohorts of 2500 or more, the difference between this and the sample mean is very small.

### 2.7 Implementation

We implemented our method in C++ using the the Eigen matrix library (Gaël Guennebaud, Benoît Jacob and others, Eigen v3, http://eigen.tuxfamily.org) for matrix manipulations and HTSlib (Li *et al*., 2009) for streaming the input VCF/BCF files.

### 2.8 Data

#### 2.8.1 Low-coverage data

We make use of two different publicly available large cohorts to evaluate our method in the low-coverage scenario. First, the 1000GP Phase 3 samples which consist of 2535 samples from a heterogeneous mix of 26 populations, each sample sequenced to an average of 7.4×. Second, data from the UK10K control group, a more homogeneous cohort than the 1000GP samples comprising 3 781 samples, each sequenced to around 7×. We only evaluated SNPs in these comparisons.

287 individuals from the 1000GP cohort were also sequenced to high-coverage (80× or more) by Complete Genomics (CG) and we used this data to validate our 1000GP calls. To create validation data for the UK10K samples, we took 63 of the European CG samples and calculated genotype likelihoods at the UK10K sites for these samples from their respective low-coverage BAM files using bcftools (Li *et al*., 2009). MarViN imputation was performed in 200kbp windows with an overlap of 100kbp between windows. We performed a number of small timing experiments on a 2Mbp region of chr20, and a more rigorous accuracy experiment using the entire chr20 for both cohorts. A summary of the samples and number of variants is in Table 1.

**Table 1.** Summary of the number of samples and SNPs for each LCWGS data set. Sample size is the number of samples present in the input genotype likelihoods, for UK10K this includes the UK10K control cohort (3 781 samples) plus an additional 63 CG validation samples with GLs calculated from low coverage alignments. Number of SNPs is the number of non-singleton bi-allelic SNPs in each respective cohort on chromosome 20. The rightmost two columns count the number of samples and SNPs that are also in the CG validation data.

| Cohort | Samples | SNPs | CG sample | CG SNPs |
| --- | --- | --- | --- | --- |
| 1000GP | 2 535 | 1 628 533 | 287 | 565 991 |
| UK10K | 3 844 | 489 278 | 63 | 223 528 |

On both these cohorts, we compared MarViN with two alternative genotype refinement schemes: Beagle 4.0 (r1399) (Browning and Browning, 2007) and the “no-LD” method we described in Section 2.1, which does not use LD information. We chose Beagle as a comparison due its popularity, ease-of-use and relative speed compared to other HMM routines (notably SHAPEIT is more accurate but also slower). Given we expect MarViN to be substantially faster, but also less accurate, than Beagle, it is reasonable to conclude that MarViN will be faster (and less accurate) than other more computationally demanding HMM based routines.

#### 2.8.2 High-coverage data

We took 50× coverage of 100bp paired reads sequenced from the widely studied NA12878 sample (ENA AC:ERR194147). These were aligned with BWAMEM 0.7.12 (Li, 2013) and small variants were called according to GATK3.3–0 best practices (DePristo *et al*., 2011; Auwera *et al*., 2013), the associated genotype likelihoods were supplied to MarViN. If an alternate allele for a variant in the 1000GP reference panel was not detected in a given sample then we used the genotype likelihood taken from the homozygous-to-reference interval in the gvcf file that overlapped the variant site.

MarViN can only improve genotyping at variants seen in the reference panel (variants with LD and frequency information). Any variant called in an individual that has been seen in a curated panel such as 1000GP is likely to be real given sufficient coverage (some amount of false discovery in 1000GP notwithstanding), since these variants have already been carefully filtered. Variants called in an individual that are not present in 1000GP require more scrutiny, although we still expect tens to hundreds of thousands of novel (mostly rare) variants in a given sample.

Hence we apply the hard filters described in Li (2015) to non-1000GP variants using hapdip (http://bit.ly/HapDip). For variants called by GATK that intersect with 1000GP, we are less stringent, only filtering on the genotype quality (GQ) field, the phred-scaled probability that a genotype is incorrect. The GQ field is produced both by GATK and MarViN.

When setting up the reference panel, we excluded NA12878 and all other CEPH1463 pedigree members from the 1000GP Phase 3 panel so as not to bias results. We only considered bi-allelic SNPs with an alternate allele count of at least five, reasoning that very rare variants were unlikely to benefit greatly from LD-based refinement. We ran MarViN for five iterations with a window size of 210kbp with overlap of 5kbp at each end (so each window overlaps by 10kbp).

As truth data, we used the highly accurate NA12878 call set from Platinum Genomes v7.0.0 (http://www.illumina.com/platinumgenomes). This consists of variants and confident homozygous-reference intervals generated from multiple aligners/callers on the 17-member CEPH1463 pedigree. The reliability of the variant calls is enhanced by retaining only those calls whose inheritance pattern across the pedigree is consistent with Mendelian inheritance. GATK/MarViN callsets were compared to this truth data using hap.py (https://github.com/Illumina/hap.py), a tool which compares variants via alignment and exact matching.

## 3 RESULTS

### 3.1 Low-coverage genotype refinement

We first evaluated each method’s speed and accuracy as a function of sample size by sampling subsets of the UK10K cohort of sizes *N* ={100, 200, 500,1000, 2000, 3844} and performing genotyping on a 2Mbp window of chromosome 20 (35Mbp to 37Mbp) containing 14416 SNPs. We measured the non-reference discordance (NRD) of each method, which is defined as (*FP+TP*)/(*FP+TP+FN*), where *TP, FP* and *FN* count the number of true positive, false positive and false negative alternate allele calls respectively. Timings were performed on a an Intel Xeon E5–2670v2 CPU with no other compute intensive processes running. We do not report compute times for no-LD as this process is dominated by I/O operations.

Figure 1 plots NRD (left) and compute time in hours (right) against sample size. When *N*=100; no-LD, MarViN and Beagle had NRD of 5.74%, 5.20% and 1.26% meaning MarViN was substantially less accurate than Beagle. However, MarViN’s accuracy dramatically increases with sample size. MarViN had 0.71% NRD at *N* = 1000 and 0.63% at *N*=3844 versus 0.59% and 0.38% for Beagle. Although still less accurate than Beagle, MarViN’s speed advantage widens with increasing *N*, it being 104× and 1445× faster than Beagle for *N*=1 000 and *N*=3 844 respectively. Notably MarViN had around 9-fold fewer errors than no-LD at *N*=3 844 and required minimal compute resources (≈14 CPU minutes for *N* = 3 844).

**Fig. 1:**
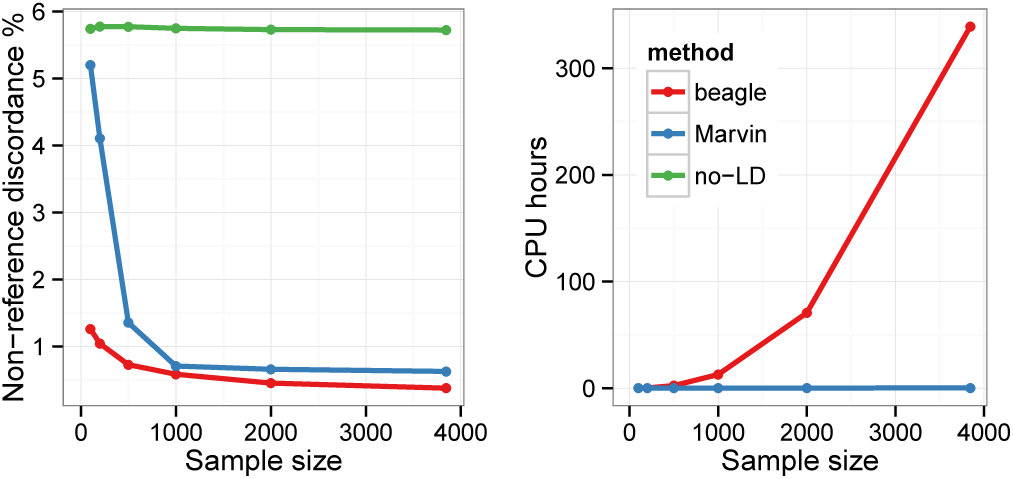
Performance of each method on UK10K cohort for increasing samples sizes (`chr20:35M-37M).` **Left:** Non-reference discordance versus sample size for three different methods. **Right:** Total compute time in hours versus sample size.

We then evaluated each method for both the 1000GP Phase 3 cohort (*N*=2 535) and the UK10K (*N*=3 844) for the entire chromosome 20 (1.63 and 0.49 million SNPs respectively). To achieve a fair comparison of compute requirements, we gave each method exclusive use of a 20-core compute node (2×10-core E5–2670v2 CPUs), running Beagle with 20 threads and running 20 simultaneous MarViN processes (concatenation time is included in the results).

Table 2 summarises the accuracy and compute times. MarViN was 360x faster than Beagle on UK10K and 46× faster on 1000GP. MarViN’s speed advantage on 1000GP is decreased relative to UK10K due to a much larger number of SNPs. MarViN had higher NRD than Beagle with 1.66% versus 0.90% on 1000GP and 0.64% versus 0.41% on UK10K. Whilst these accuracy differences may seem small, the error rates are concentrated at low frequency genotypes. Nevertheless, MarViN has 4.64× and 9.82× fewer errors than the naive no-LD routine.

**Table 2.**
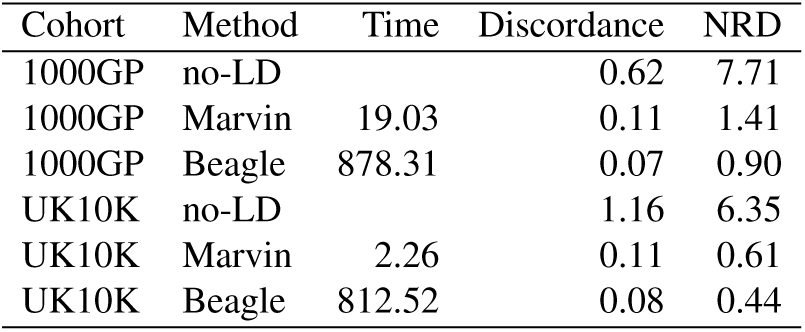
Performance for each method on the 1000GP (*N* = 2535) and UK10K (N = 3844) chr20 data. Time is the compute time in hours on a 20-core (2×E5–2670v2 CPUs) server with 132GB RAM when using 20 threads. Discordance is the percentage of discordant genotypes between the imputed genotypes and high-coverage genotypes on the CG validation samples. Non-reference discordance (NRD) is the discordance when not counting genotypes that were homozygous reference in both the imputed and high-coverage genotypes.

We then investigated accuracy at different allele frequencies by binning genotypes by allele frequency and calculating Pearson’s correlation coefficient (*r*^2^) between the imputed genotypes and the high-coverage validation genotypes within each bin. Figure 2 plots *r*^2^ against allele frequency (log 10 scale) for 1000GP (left) and UK10K (right). We see for common variants (AF≥2%) Beagle and MarViN are roughly equal (and substantially better than no-LD). Beagle outperforms MarViN at lower allele frequencies. Figure 2 and Table 2 also suggest that MarViN performs less well on heterogeneous cohorts such as 1000GP, compared to relatively homogeneous cohorts like UK10K.

**Fig. 2:**
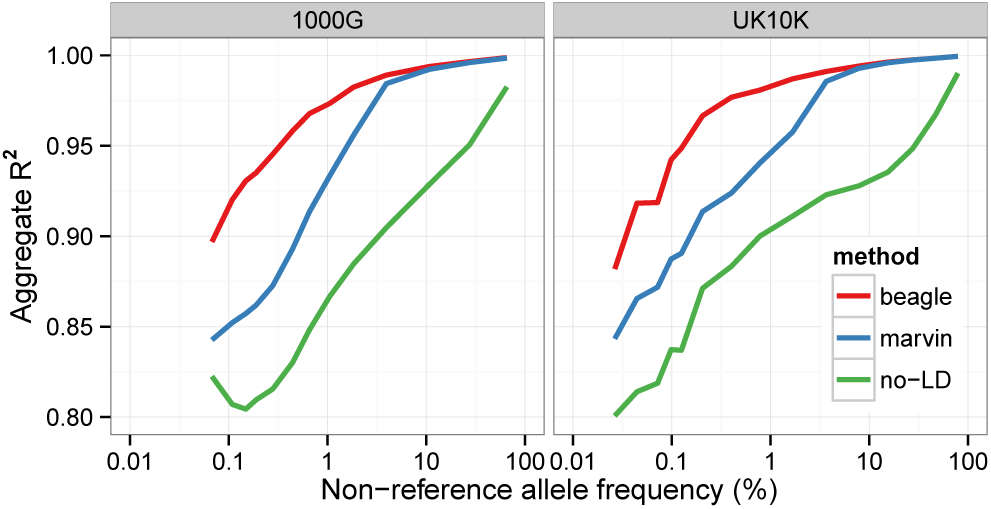
Pearson’s correlation coefficient between imputed and true genotypes for different cohorts as a function of allele frequency for different data sets. **Left:** 1000GP **Right:** UK10K.

### 3.2 High-coverage genotype refinement

Figure 3 plots recall (proportion of PG SNPs detected and correctly genotyped) against precision (the proportion of called SNPs that are concordant with PG) for GATK before and after refinement with MarViN for increasingly liberal filters on the GQ field. MarViN refinement yields a modest, but consistent, improvement in SNP recall for a given level of precision (and vice versa).

**Fig. 3:**
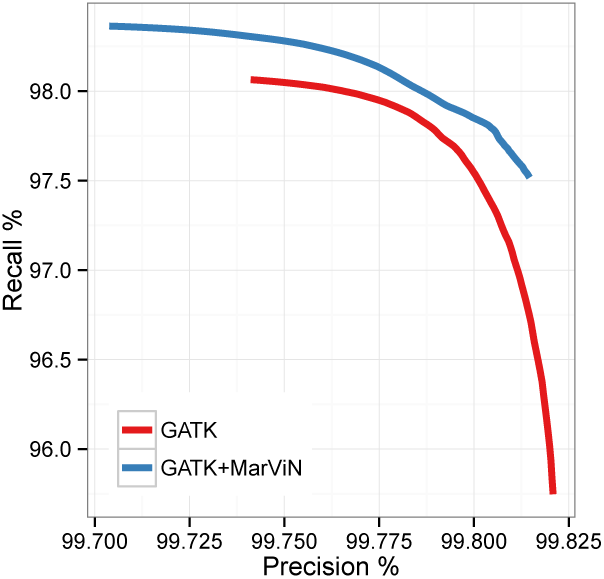
Recall versus precision for GATK SNP genotype calls before and after refinement with MarViN for increasingly aggressive GQ filters. Recall is defined as the proportion of PG alternate genotypes correctly identified. Precision is the proportion of alternate genotypes called that are concordant with the PG data set.

Table 3 further breaks down these results. Firstly, there were 243 381 SNPs called by GATK that were not in 1000GP (with minor allele count >4), these were filtered using the hard filters in hapdip. Such SNPs cannot be further refined but we report them for completeness. Of these, 143 247 SNPs were validated in the Platinum Genomes dataset and a total of 2 386 were classified as false positives due to having either incorrect genotypes, incorrect alleles or being called in a known homozygous-reference region. This yields a precision of 98.36%, which as one might expect, is lower than calls that intersect with 1000GP variants.

**Table 3.** Summary of high coverage SNP calls by GATK that were not present in 1000G with MAC greater than 4 (first column), calls made by GATK that were present in 1000G (middle column) and GATK calls after applying Marvin genotype refinement (third column).

|  | non-1000G | 1000G |  |
| --- | --- | --- | --- |
|  |  | GATK | Marvin |
| Total | 243 381 | 3 387 126 | 3 408 128 |
| Unevaluated | 97 748 | 74 813 | 86 845 |
| TP | 143 247 | 3 309 226 | 3 317 953 |
| FN | 3 377 480 | 211 373 | 202 888 |
| FP1 (GT wrong) | 1 251 | 1 577 | 1 318 |
| FP2 (allele wrong) | 98 | 81 | 96 |
| FP3 (homref) | 1 037 | 1 429 | 1 916 |
| FP (total) | 2 386 | 3 087 | 3 330 |

For SNP calls that intersect 1000GP, we only applied a GQ≥20 filter. The GATK callset contained 3 387 126 SNPs, rising to 3 408 128 after refinement with MarViN. Of these, 3 309 226 and 3 317 953 were validated as correct in Platinum Genomes, meaning MarViN refinement yielded an additional 8 727 correctly genotyped SNPs. In terms of effect on false positive rate, MarViN reduced the number of incorrect genotypes (with correct allele) from 1 577 to 1 318, as one might expect *genotype refinement* to do. However, Marvin also imputed a greater number of variants with incorrect alternate allele (96 versus 81) and SNPs in homozygous reference regions (1 916 versus 1 037). This means MarViN had a slightly higher number of false positives than the raw GATK callset, 3 330 versus 3 087, bringing its precision to 99.902% versus 99.909% for GATK 99.902%. Given the gains in SNP recall, this seems a minor cost to pay.

## 4 DISCUSSION

The algorithm presented in this paper is at least two orders of magnitude faster than Beagle on the UK10K cohort. Whilst this speed does come with a decrease in accuracy (particularly for rare variants), our method still makes nearly 10-fold fewer errors than a genotyping routine that does not take linkage-disequilibrium into account.

The rapidly growing size of reference panels may soon preclude the use of super-linear complexity techniques such as Beagle, since computation will become too expensive. For example, the Haplotype Reference Consortium (http://www.haplotype-reference-consortium.org) has collected around 35 000 LCWGS samples to create a reference panel for imputation. Extrapolating from Figure 1, it seems unlikely it would be tractable to run Beagle on a cohort of this size. One possible use of MarViN would be to quickly generate an initial estimate of genotypes, which could then be supplied as starting values to a more sophisticated routine, reducing the number of iterations the latter needs to perform. MarViN might also be an ideal routine for intermediate coverage (≈15×) projects.

The reduced accuracy of MarViN compared to Beagle at lower frequency variation is likely due to the limitations of modelling the population using one vector of allele frequencies and one covariance matrix. This simplistic model may not capture more subtle population substructure. Notably MarViN performs better on the more homogeneous UK10K cohort than on the 1000GP cohort which has far more population structure (although also has a smaller sample size). One possible way to improve this situation would be to add more flexibility to the MarViN model by using an MVN mixture distribution, but we leave this for future work.

We have also demonstrated the efficacy of genotype refinement in the high-coverage scenario, the first such investigation to our knowledge. A modest gain in recall for SNPs was achieved at a cost of a negligible decrease in precision. We also attempted refining indels with this approach, gains in recall were indeed observed but were accompanied by unacceptable increases in the false-discovery rate. This may be due to a higher FDR in the 1000GP indels and could perhaps be solved via aggressive filtering.

While the improvements seen on high-coverage data are modest, we nevertheless believe it noteworthy that results achieved from high-coverage data can be improved at all by this method. Moreover the efficiency of our method means it adds little additional overhead to processing pipelines for WGS data, whereas genotype refinement using existing HMM-based methods would be a considerable computational undertaking.

## ACKNOWLEDGEMENTS

We thank Stathis Kanterakis for providing GATK results.*Funding*: All authors are employees of Illumina Cambridge Ltd., a public company that develops and markets systems for genetic analysis, and receive shares as part of their compensation.

